# Parsing the roles of neck-linker docking and tethered head diffusion in the stepping dynamics of kinesin

**DOI:** 10.1101/183590

**Authors:** Zhechun Zhang, Yonathan Goldtzvik, D. Thirumalai

## Abstract

Kinesin walks processively on microtubules (MTs) in an asymmetric hand-over-hand manner consuming one ATP molecule per 16 nm step. The contributions due to docking of the approximately thirteen residue neck linker to the leading head (deemed to be the power stroke), and diffusion of the trailing head contribute in propelling the motor by 16 nm have not been quantified. We use molecular simulations by creating a new coarse-grained model of the microtubule-kinesin complex, which reproduces the measured stall force as well as the force required to dislodge the motor head from the MT, to show that nearly three quarters of the step occurs by bidirectional stochastic motion of the TH. However, docking of the neck linker to the leading head constrains the extent of diffusion and minimizes the probability that kinesin takes side steps implying that both the events are necessary in the motility of kinesin, and for the maintenance of processivity. Surprisingly, we find that during a single step the trailing head stochastically hops multiple times between the geometrically accessible neighboring sites on the MT prior to forming a stable interaction with the target binding site with correct orientation between the motor head and the *α/β* tubulin dimer.

**Significance Statement:** Like all motors, the stepping of the two headed conventional Kinesin on the microtubule is facilitated by conformational changes in the motor domain upon ATP binding and hydrolysis. Numerous experiments have revealed that docking of the thirteen residue neck linker (NL) to the motor domain of the leading plays a critical role in propelling the trailing head towards the plus end of the microtubule by nearly 16 nm in a single step. Surprisingly our molecular simulations reveal that nearly three quarters of the step occurs by stochastic diffusion of the trailing head. Docking of the NL restricts the extent of diffusion, thus forcing the motor to walk with overwhelming probability on a single protofilament of the MT.

---

Directional transport of intracellular vesicles along polar tracks (actin and microtubule (MT)) is carried out by a variety of molecular motors (1). The crucial function is executed by the three families of motor proteins (myosin, kinesin, and dynein), many of which preferentially move towards a certain direction along a particular polar cytoskeletal track (2). For example, kinesin-1 (Kin1) or conventional kinesin, pulls cargo towards the plus (+) end of the microtubule (MT) (2-4). It is now firmly established, thanks to a number of high precision experiments, that kinesin with two motor domains walks in a hand-over-hand manner (5-8), by consuming one ATP molecule per step (9). Despite the small size of the motor domain, Kin-1 is a powerful and fast motor, moving towards the (+) end of the microtubule (MT) resisting forces up to 7 pN (10, 11), which is larger or equal than the stall forces of bigger motors (12, 13), such as dynein (1-7) pN and myosin (approximately 3 pN). Kin-1 moves towards the (+) end processively at a speed of ~ 800 nm / sec (10), which is greater than both myosin V (~ 400 nm/s) and dynein (~ 100 nm / sec) (14).

A remarkable series of experimental studies (3, 6, 15-17) from a number of groups has revealed many of the details of the stepping mechanism of kinesin. Two mechanisms have been proposed to explain how kinesin converts chemical energy to mechanical work to walk towards the (+) end of the MT in a hand-over-hand manner. According to the “power stroke” model (8), neck linker (NL) docking induced by ATP binding to the microtubule-bound leading head (LH), pulls the trailing head (TH) into the neighborhood of the target binding site (TBS) that is 16 nm away from the initial binding site. In this model, the NL with ~ 13 residues connecting the motor domain to the coiled coil, may be structurally analogous to the easily identifiable lever arm in myosin motors (2). In contrast, “Brownian ratchet” model (18) posits that ATP hydrolysis in the TH allows it to detach from microtubule to initiate biased diffusional search towards the TBS.

Both models have experimental support. Experiments using single molecule FRET (Fluorescence Resonance Energy Transfer) (19) and fluorescence anisotropy (20) show that the neck linker docks (power stroke) upon ATP binding to the LH, raising the possibility that the motility of kinesin arises predominately from the power stroke mechanism. An optical trap experiment (21) shows that a kinesin mutant, which lacks the cover strand (a major docking site of the NL), can still walk processively but generates much less force. It can, therefore, be concluded that neck linker docking must contribute significantly to force generation. In all likelihood, both power stroke and diffusion are operative in the stepping of kinesin. Until recently (22) the extent of diffusive motion of kinesin has not been reported in experiments (3), although a number of observations support the importance of the Brownian ratchet model. First, the small size of even a fully stretched NL limits the potential physical displacement of the TH upon NL docking to the LH (3). Second, the temperature dependence of the stepping rates indicates an entropic nature of directional bias (18) that cannot be explained solely by the

NL docking model. However, the observation that the mutant lacking cover strand walks processively indicates that kinesin could walk by a Brownian ratchet mechanism in the absence of external load.

The two mechanisms are not mutually exclusive (3, 23, 24). Therefore, an unresolved question is what fraction of the kinesin step is associated with power stroke and diffusion, respectively (3, 25)? If a large fraction of the step is associated with power stroke, we expect that the TH to move almost unidirectionally covering a majority of the 16 nm, and the bidirectional diffusion could occur only within the neighborhood of the TBS. On the other hand, if a majority of the 16 nm step is covered by diffusion of the TH, we expect that the motion of the TH to be bidirectional, and the extent of stochastic random walk of the TH to be large (> 8 nm, half of the step size). However, if the kinesin step is largely diffusive, what keeps the motor on a single protofilament of the microtubule (26-28), which contains multiple protofilaments?

In order to distinguish between the predictions of these two (extreme) models the motion of the kinesin motor head has to be tracked at a microsecond temporal resolution. Despite spectacular advances in using microscopy methods (see for example (22)) to visualize the stepping kinetics of motors, the needed time resolution to unambiguously track the position of the stepping motor has not been reached. Numerous experiments, starting with the pioneering study by Block and coworkers (29), which map the motion of the kinesin head (or the position of the load) at lower temporal resolution show an apparently unidirectional step between the initial binding site (0 nm) and the TBS (16 nm). The mechanism of TH motion is “hidden” in the jump time (~ 30 ***μs)*** between the waiting state, detachment of the TH, and subsequent attachment to the MT (both heads are bound to the MT) of kinesin. Unless experiments can track the molecular events in the motor head on shorter time scales (~ (5-10) μs) one cannot unambiguously assess the interplay of power stroke (involving a structural change in the neck linker) and stochastic motion of the tethered head in resolving the 8 nm step of kinesin.

Given that a globular protein of the size of kinesin head with radius of gyration ***R_g_*** ≈ 2 nm, can diffuse 16 nm within 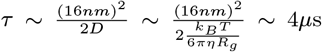 in a medium as viscous as cytoplasm, microsecond resolution may be needed to capture the potential importance of the stochastic bidirectional motion of the TH. However, due to difficulties in tracking kinesin experimentally at microsecond resolution (3), the importance of diffusivity of the kinesin step has been stressed (3, 18) but not been quantified.

Here, we use Brownian dynamics of a coarse-grained (CG) model of microtubule-kinesin (MT-Kin) complex to monitor the motion of kinesin during the 16 nm step at high temporal resolution (24). Such models (30) have provided considerable insights into a variety of complex biological systems, as first illustrated by Hyeon and Onuchic (HO) (23) in the context of stepping of kinesin. Our simulations, which produce physically realistic picture of how kinesin steps on MT, allow us to follow the motion of the TH and the LH neck linker separately at sub-microsecond resolution by generating several hundred trajectories. We show that a substantial portion of the kinesin step occurs by a diffusive process. However, NL docking provides severe restrictions on the conformational space explored by the TH during the stochastic search for the TBS. Thus, a combination of NL docking and diffusive search for the TBS (16 nm away) is needed for executing the movement of the TH predominantly towards the (+) end of the MT in the absence an external resistive force.

## Results

Our model, which mimics the typical single molecule experimental set up closely, consists of the two heads bound to the MT in the resting state. Following our previous study (24), we include three MT protofilaments, two motor heads, a coiled coil (length ≈ 30 nm) and a 500 nm spherical cargo (Fig. S1 in the Supplementary Information (SI)). The coiled-coil is connected to the junction at which the two neck linkers, one from each motor domain, meet. The spherical cargo is attached to the ends of the coiled-coil. In order to probe the possibility that the detached motor head could explore the binding sites on the neighboring protofilaments we created a model (details in the SI) containing three of the thirteen MT protofilaments.

### Two energy scales determine kinesin motility

In order to ensure that the simulations are realistic, we first reproduced two important experimentally measured mechanical properties of the kinesin motor, the stall force (F_s_) (10, 11) and the force required to unbind TH from the MT (***F_u_***) (31). We expect that ***F**_s_* and ***F**_u_* should depend on the two energy scales 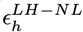 (the interaction strength between the NL and the LH) and 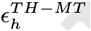 (the interaction between the TH and the MT) (24). Determination of the range of 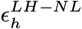 values that reproduces the measured ***F**_s_* is needed for a realistic description of the NL-LH interaction, which largely determines the role of power stroke in facilitating the kinesin step. Furthermore, the model has to reproduce the measured value of ***F**_u_* that depends on realistic modeling of the TH-MT interaction, which in turn affects not only the probability that Kin1 could take side steps but also determines the final stages of motor head-MT recognition (24). The two energy scales, needed to reproduce ***F**_s_* and *F_u_*, are independent. Residues participating in the LH-NL interaction are in the leading head (LH). In contrast, residues essential for TH-MT interaction are in the trailing head (TH). In addition, our previous study [24] demonstrated that 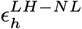 and 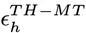 affect different stages of kinesin step.

### Determination of

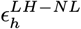 For each 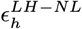, we first performed a set of control simulations in which no external force is applied to kinesin (black curve in Fig. S2b), and then another set of simulations in the presence of a resistive force of 7 pN (Fig. S1a and blue curve in Fig. S2b). For each set, we measured the probability (***P**_f_*) that TH steps forward to the TBS, as well as the probability (***P**_b_*) that TH goes back to the initial binding site (IBS). Optical trap experiments (10, 11) indicate that, ***P_f_ = P_b_*** at the stall force, F_s_=7pN. Our simulations show that only for a narrow range of 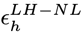, ***P_f_ ≈ P_b_*** at ***F**_s_* = 7pN. At ***F**_s_*, with 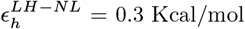, we find that ***P_f_ ≈ P_b_***. (Figs. S2b-S2c). In a control simulation with F = 0, kinesin predominately moves forward (Figs. S2b-S2c). In the rest of this paper we set 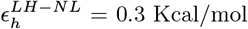.

### Calibrating the MT-TH interaction by reproducing the unbinding force

(*F_u_*). Experiments show that the unbinding force for monomeric kinesin varies from 3 pN to 9 pN depending on the nucleotide condition (31). In order to obtain 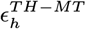, We performed simulations by initially binding a single motor head to the MT (Fig. S2d), just like in the experimental set up, to obtain ***F**_u_*. By performing hundreds of simulations using various combination of 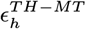 and ***F***, we were able to find the values of 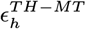 that allow kinesin to bind stably to the MT in the absence of F (black curve in Figs. S2e-2f) but detaches at 3 pN corresponding to the weakly bound state (blue curve in Fig. S1e) or 9 pN corresponding to the strongly bound state (blue curve in Fig. S2f). Therefore, by varying 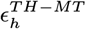 we can mimic both the weak and the strong binding states that kinesin experiences during the reaction cycle. Because the exact timing and condition for ADP release, which strengthens the MT-Kin interaction is still unknown, here we perform simulations under conditions that mimic both weak and strong binding.

### Translation motion of the trailing head is diffusive

A key finding in our simulations is that the search for the TBS is a predominately diffusive process, independent of our choice of 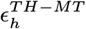. We observe large scale bidirectional diffusive motion of the TH throughout the 16 nm step. For example, four representative trajectories where TH completes the 16 nm step (Fig. 1a-1d), with varying first passage times, show that the center of mass of the TH fluctuates extensively (along the MT axis). The duration of such bidirectional diffusional search varies from < 10*μs* (Fig. 1a) to > 100*μs* (Fig. 1d).

**Fig. 1.**
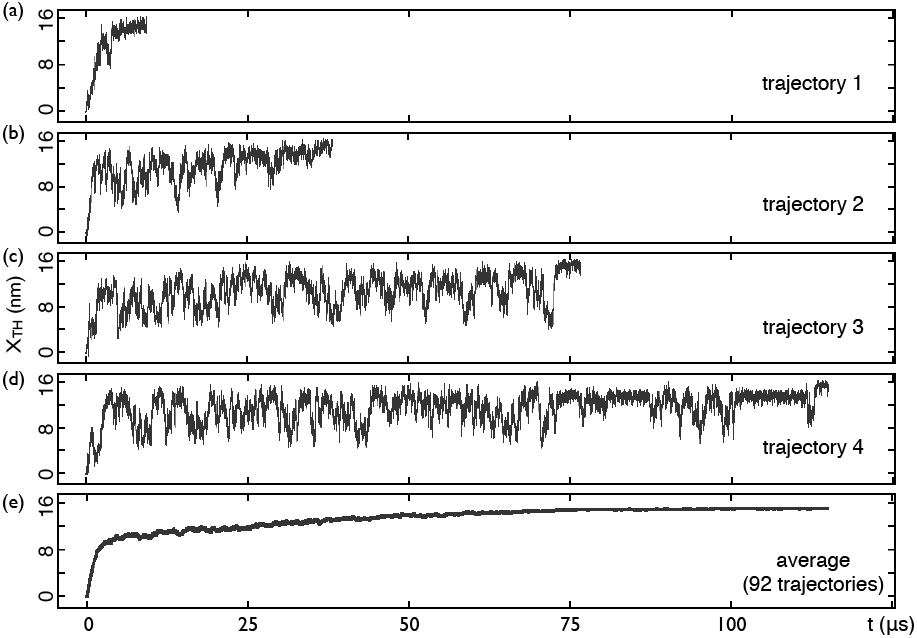
Diffusive nature of the kinesin step: (a-d) Translational motion of (the center of mass) of the **TH** along the **MT** axis in four trajectories of the 16 nm step of kinesin, quantified using the time dependent changes of the center of mass of the trailing head, *x_TH_*, along the **MT** axis. (e) Ensemble average of *x_TH_* based on 92 trajectories. The average can be best fit using *x_TH_* (t) = 15.7 — 8.5e^(-*t*/1.1*μs*)^ — 6.4e^(-*t*/40.1*μs*)^, where the coefficient are in units of nm. Note that at *t* = 0, *x_TH_* (*t* = 0) = 0.9nm, which is roughly the equilibrium value of *x_TH_*. At long times *x_TH_* = 15.7nm. Thus, *x_TH_(t)* increases from *t* = 0 till stepping is completed.

The ensemble average of *x_TH_*(*t*) (Fig. 1e) also shows that the **TH** spends considerable amount of time undergoing stochastic motion even as it is poised to reach the **TBS**. The average motion of the **TH** (along the **MT** axis, based on 92 trajectories) shows a fast and slow phase. The fast phase occurring within 1.1 ***μs***, corresponds to the relaxation of the TH after detachment from the **MT** and the forward motion induced by neck linker docking in the **LH**. The slow phase with an average time constant of 40.2 μs, corresponds to the diffusional search for the TBS. We find that the fast and slow phases are robust to choice of 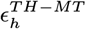 (see Fig. S3). The time constant for the slow phase does not significantly depend on 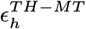 as long as its value is in a range for stable binding of the **TH** to the **MT**. However, if 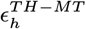 is too weak the TH could spontaneously detach due to thermal fluctuation and motor processivity would be lost. As 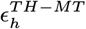 decreases, the time constant corresponding to the slow phase could exceed 40.2 μs, an estimate based on a value of 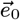 value that reproduced the experimental unbinding force of (6-9) pN. From the additional simulations and the arguments given above we conclude that for physically reasonable values of 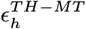 the time constants are not greatly affected, allowing us to conclude strongly that temporally more than 95% of the kinesin step is associated with diffusion.

A close examination of a typical trajectory (Fig. 2) reveals that diffusion starts immediately after the **TH** detaches from the **MT**. Even when ***x_TH_*** reaches 10 nm at ~ 1 *μ*s, it subsequently decreases to ~ 3 nm (Fig. 2e). At a later time the TH enters the neighborhood of the TBS at ***t ~*** 17 *μ*s (see Fig. 2b and the arrow in Fig. 2f), but again retreats to a position behind the MT-bound leading head at ~ 20 *μ*s (Fig. 2e). Additional evidence for diffusion outside the neighborhood of the TBS comes from the recording of ***d**_TH_*, the distance between the **TH** and the **TBS**, as a function of time (Figs. 2c and 2f). It is also clear from Fig. 2f that the **TH** stochastically searches for the **TBS** upon detachment from the **MT**. The diffusive characteristics of the trajectory depicted in Fig. 2 is typical, and we find similar behavior in all the other stepping trajectories as well (for example the ones illustrates in Fig. 1). We surmise from the time-dependent changes in both ***x_TH_*** and ***d_TH_*** that the TH undergoes bidirectional diffusion not only within the neighborhood of the TBS, but also throughout the 16 nm step.

**Fig. 2.**
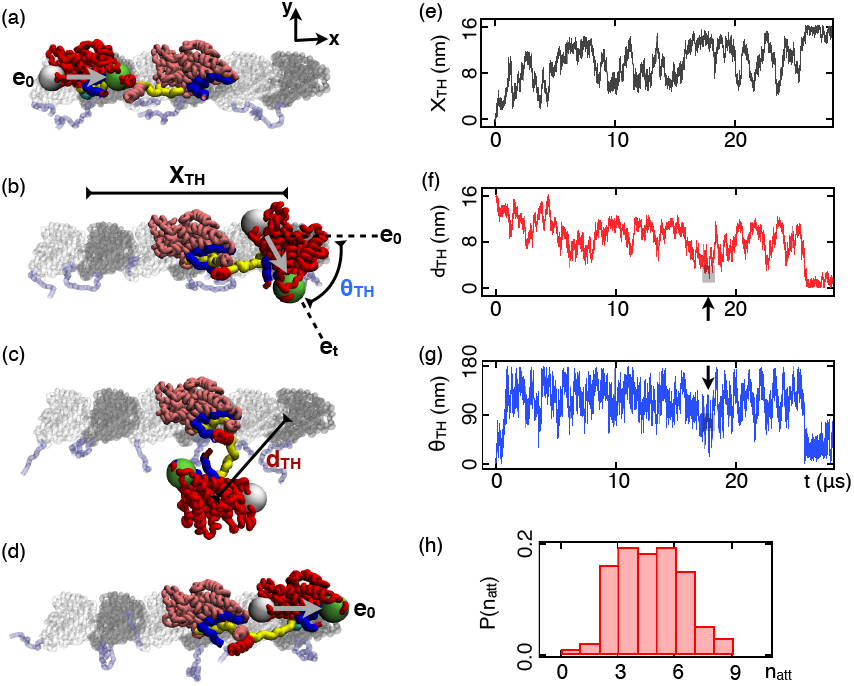
Stochasticity in a 16 nm step of kinesin: (a-d) Four snapshots (at 0.0, 17.1, 19.3, and 27.8 *μ*s) in a representative trajectory of the 16 nm step of kinesin. The trailing head (TH) is in red, and the leading head (LH) is shown in pink. Yellow structure is the neck linker, and the docking site for the NL is in blue. a and /3-tubulin are in silver and grey, respectively, and are augmented by ehooks (light blue). The black arrows (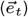) indicates the orientation of the **TH** during the step, where 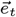 is an unit vector pointing from residue V40 to residue N221 of the TH. (e) Record of the translational motion of *x_TH_*, the center of mass of the **TH**, along the **MT** axis during the 16 nm step. (f) The dependence of d the distance between the **TH** and the **TBS** as a function of t, in a representative trajectory. One unsuccessful attempt **TH** made to bind to the **TBS** is highlighted by the black arrow. (g) Time-dependent changes in the rotational motion, quantified using *θ_TH_* (t), of the **TH** (with respect to its center of mass) during the same 16 nm step as (b). (h) Distribution of n*_att_*, the number of times the **TH** reaches the **TBS** with incorrect orientation (the number of unsuccessful attempts), based on 92 trajectories.

### Trailing head undergoes isotropic rotational diffusion

Time-dependent changes in ***x_TH_*** and ***d_TH_*** reveal only one facet of diffusive behavior of the **TH** during the kinesin step. The **TH** also undergoes rotational diffusion. We use *θ**_TH_*** (Fig. 2b) to quantify the extent of rotation of the **TH** with respect to its center of mass. At t = 0, ***θ_TH_*** ≈ 0° (Fig. 2g) implying that the **TH** is bound to the MT with the same orientation as observed in CryoEM image of the kinesin-MT complex (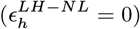 in Fig. 2a). We assume that stepping is complete only after the **TH** achieves the same orientation in the **TBS** (***θ_TH_*** ≈ 0°). During the stepping process, *θ**_TH_*** changes randomly between 0° and 180° (Fig. 2g). In the representative trajectory shown in Fig. 2, the **TH** is in the vicinity of the **TBS** at ~ 17.5 μs. However, at this instant the value of *θ**_TH_*** is close to 70° (Fig. 2g), which implies that one of the principal axes of the **TH** that is initially parallel to the MT axis when the **TH** is bound to the **MT** (see the black arrow in Fig. 2a), is almost perpendicular to the **MT** axis (see the black arrow in Fig. 2b). Because of the incorrect orientation, the **TH** fails to bind to the **TBS** at 17 μs, and diffuses away from the TBS. Only at ***t ~*** 26 ***μs*** does the **TH** achieve the correct orientation (~ 0°, see Fig. 2d).

The rotational motion of the **TH** is as important as translation, because kinesin head cannot bind to the **MT** and function with incorrect orientation (***θ_TH_*** ≠ 0). It has been shown, using alanine scanning, that all residues responsible for **MT** binding are located on one side of kinesin (32). Furthermore, crystal structures of the intermediate states during Mg-ADP release and CryoEM structure of the MT-Kin complex suggest that activation of kinesin requires multiple specific contacts with the **MT** (33). These results imply that stable binding between the **TH** and **MT** as well as the function of kinesin requires specific orientation between the motor head and the **MT**.

To further demonstrate the importance of rotational diffusion of the **TH**, we calculated ***n_att_*** the number of times the **TH** retreats from the **TBS** due to incorrect orientation (Fig. 2h). In the trajectory shown in Fig. 2g, ***d_TH_*** transiently reaches ~ 0 nm at ~ 17 μs. However, it remains unbound, at t~ 20*μ*s because *θ_TH_* ≠ 0. The distribution of the number of such failed attempts (Fig. 2h), based on 92 trajectories, shows that only in less than 5 % of trajectories the **TH** binds to the **TBS** with the correct orientation at the instant when ***d_TH_ ~*** 0. Typically, the **TH** lands and unbinds ~(3-6) times before it can finish the 16 nm step that satisfies both the distance (***d_TH_ ≈*** 0) and orientational (***0_TH_ ~*** 0°) criteria. This explains why in many trajectories the 16nm step takes more than 30 ***μ*** s to complete, even though the first passage times estimated based on translational diffusion coefficient and rotational diffusion coefficient (see Fig. S4) are only 3.0 μs and 4.0 ***μ*** s, respectively. In other words, the first passage time is not the actual binding time (34). We note here in passing that neither the translational nor the rotational diffusion depends strongly on 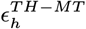 (see Fig. S4). Our simulations show that a great deal of stochasticity is involved in achieving an interface between the **TH** and **MT** that satisfies both the distance and orientation criteria. Because of the diffusive nature of the motion, once the **TH** leaves the TBS, it could take the **TH** more than 10 *μs* to return to the **TBS** (between 17 *μs* and 25 *μs* in Fig. 2g for example). During this time interval, the **TH** may diffuse as far as 10-12 nm away from the TBS, as illustrated in the time-dependent change in ***d(t***) after 17 *μs* in Fig. 2f. Taken together, our results show that search for the **TBS** involves both translational and rotational diffusion. Rotational diffusion of the **TH** is isotropic but the anisotropic translation motion is greatly (in the absence of applied resistive force) biased towards the (+) end of the **MT** (24), which, we show below, is achieved by the NL docking to the LH.

### A large diffusion length quantifies stochasticity

Our simulations allow us to quantify the fraction of the kinesin step associated with the power stroke and that due to tethered diffusion. Although such a quantification has been reported for myosin motors (35) in experiments, a similar parsing of the kinesin step is undocumented. We use hundreds of trajectories to measure the fraction associated with power stroke and diffusion. For each trajectory, we first identify the instant when the power stroke associated with neck linker docking is complete (see the inset in Fig. 3a). We calculated the diffusion length, *x_df_*, by measuring the upper and lower values of *x_TH_* that are reached after the completion of NL docking. For the trajectory in Fig. 3a, *x_TH_* fluctuates between 4 nm and 16 nm after NL docks, and therefore *x_df_* = 12 nm. The reason *x_TH_* does not fluctuate between 0 and 16 nm, is that once NL docks to the LH the limited length of the stretched **TH** neck linker prevents the **TH** from diffusing back to the initial binding site. Thus, the lower bound, 4 nm, corresponds to the fraction of 16 nm due to power stroke (***x_ps_***) in this trajectory. The distribution of ***x_ps_***, based on 100 trajectories, shows that in general the power stroke is responsible for 3-5 nm of the total 16 nm step (Fig. 3a). Therefore, the diffusion length has to be in the range ~ (11-13) nm in most of the trajectories for successful completion of a step.

**Fig. 3.**
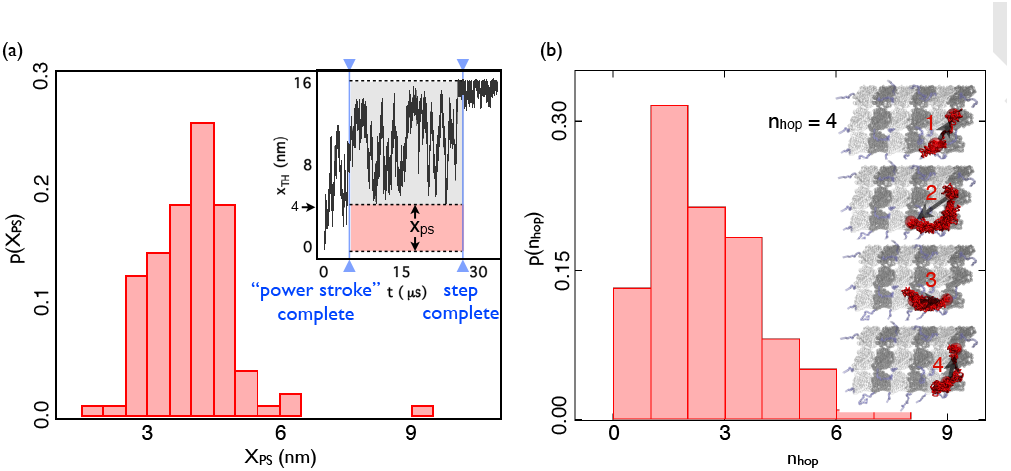
Fraction of 16 nm step associated with power stroke: (a) Distribution of *x_ps_*, the fraction of 16 nm associated with power stroke, based on 100 trajectories. The inset illustrates *x_ps_* and also shows the time variation in the translational motion of *x_TH_* the **TH** along the **MT** axis in a representative trajectory. The first vertical line (solid blue) shows the instant neck linker docking (power stroke) is complete, after which the **TH** undergoes diffusive motion. The top and middle horizontal lines (dotted blue lines) indicate the extent of diffusional search after **NL** docking. The distance between the top and middle line corresponds to the fraction associated with diffusion, while the distance between the middle and bottom line corresponds to the fraction associated with power stroke. (b) Distribution of *n_hop_*, the number of hopping events (see text for definition) within the first 30 *μ*s, based on 100 trajectories. Four hopping events in a single trajectory are illustrated on the right.

### Kinesin hops stochastically between multiple geometrically accessible binding sites on the MT

Does **TH** visit binding sites on neighboring protofilaments? This question is pertinent not only because the **MT** has multiple protofilaments but also because of our finding that nearly three quarters of the 16 nm step is associated with diffusion of the TH. We find that the **TH** not only visits neighboring protofilaments but also hops repeatedly between the binding sites on neighboring protofilaments and the TBS. For example, in a single trajectory (see the inset of Fig. 3b) the **TH** first hops from the lower right binding site to the TBS, then diffuses from the **TBS** to the binding site below the site occupied by the LH. Subsequently, the **TH** revisits the lower right binding site, before finally being captured by the TBS. On an average, **TH** hops 2-3 times within 30 ***μ***s (Fig. 3b). The average hopping rate, calculated based on hundreds of such events, is ~ 10 *μ*s^-1^.

The results in Fig. 3b suggest that the **TH** may hop multiple times during a single kinesin step, depending on the affinity between ADP-bound **TH** and the MT. If the affinity is sufficient to trap the ADP-bound **TH** at a specific binding site, which requires the correct orientation of the motor head (most likely the TBS) with respect to the **MT** the **TH** may only hop between the geometrically allowed sites on the neighboring protofilaments 1-3 times within a step. On the other hand, if none of the accessible binding sites can trap the ADP-bound **TH**, hopping of the **TH** may persist until ADP release occurs. Release of ADP strengthens MT-kinesin interaction and hence would result in the cessation of the diffusional search and stochastic hopping. Given that the ADP release time (~ ms) is much slower than the average hopping time between binding sites (~ 10 μs), it is likely that **TH** could potentially hop hundreds of times within a single step.

### NL docking constrains diffusion of the **TH** to minimize side steps

So far, we have provided four lines of evidence support the diffusive nature of the kinesin step: (i) high resolution recording of the translational and rotational motion of the **TH** (Figs. 2a-2g), (ii) multiple attempts to bind to the **TBS** (Fig. 2h), (iii) large diffusion length (Fig. 3a), and (iv) stochastic hopping between binding sites (Fig. 3b). These findings might give the erroneous impression that tethered diffusion alone could lead to a site 16 nm away on the same protofilament as the **TBS** without NL docking playing a significant role. In order to explore if this is indeed the case, we performed a “mutation” simulation, in which NL docking is energetically unfavorable (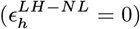). In this case **TH** is more likely to visit binding sites on neighboring protofilaments besides the **TBS** (Fig. 4a). Just as in wild type (WT) simulations (where docking is favorable), we found that the **TH** stochastically hops between the accessible binding sites due to the diffusive nature of head motion in the mutant simulations. However, in the absence of NL docking, the probability of **TH** hopping to binding sites on the neighboring protofilaments of the **MT** is much larger than the probability of reaching the TBS. In contrast, in the WT simulations, the probability of **TH** reaching the side and initial binding sites is substantially less than finding the TBS. The NL docking essentially prevents the tethered head from reaching the initial binding site, implying that unlike in Myosin V (36) rear foot stomping does not occur. Thus, the restriction imposed by NL docking facilitates kinesin to reach the **TBS**, even though the movement of the **TH** after detachment from **MT** is stochastic.

**Fig. 4.**
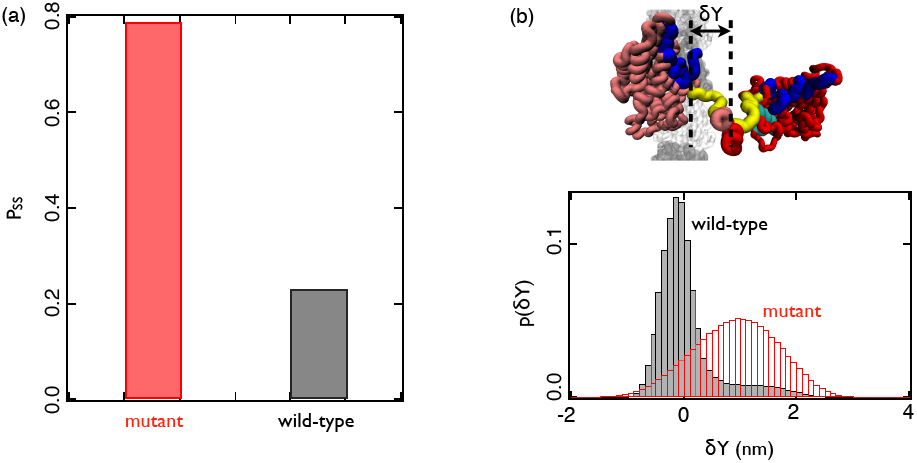
Neck linker docking decreases the probability of **TH** taking side steps: (a) Comparison of the probability of side steps in the mutant (docking is not energetically favorable) and wild type (docking is energetically favorable) simulations. Here 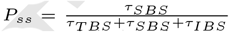 τ_SBS_(τ_SBS_ and τ_IBS_)is the average time the **TH** spends in the neighborhood of side binding sites on neighboring protofilaments (the **TBS** and the **IBS**) over 100 trajectories. The **TH** is considered to be in the neighborhood if *d* <4nm, where d is defined in the methods. (b) Distribution of sideway extension of LH-NL (δY) in the mutant and wild type simulation, where 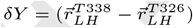 · 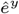 with 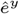 being the unit vector along the y axis (see Fig. 2).

Neck linker docking also decreases the probability of **TH** visiting side binding sites because it constrains the sideways extension of the LH neck linker. Our previous study (24) showed that in order for the **TH** to take a side step, not only the **TH** neck linker but also the LH neck linker needs to extend sideways. If docking is not energetically favorable, LH neck linker can extend sideways freely (red bins in Fig. 4b). However, the interaction between the catalytic core and the LH neck linker would limit the sideways extension of the LH neck linker if docking is energetically favorable (black bins in Fig. 4b). This in turn will reduce the diffusion coefficient somewhat (see Fig. S5) and decrease the distribution width of the **TH** along the y axis (Fig. 4b). Therefore, although NL docking contributes only 3-5 nm out of 16 nm, it plays a crucial role in restricting the movement of kinesin on a single protofilament.

### Dynamics of NL docking and the diffusing **TH** are uncorre lated

In order to illustrate how NL docking constrains the diffusion of the **TH**, we compare the displacement of the docking LH-NL and moving **TH** in a representative trajectory (Fig. S3a). Specifically, we show the motion of T338 (the red curve in Fig. S3b) at the boundary between the coiled coil and the LH-NL (Fig. S3a), which docks upon ATP binding. In the same figure, we also plot the motion of the center of mass of the **TH** (the black curve in Fig. S3b). Interestingly, the motion of the **TH** and the docking NL seems to be largely uncorrelated, except at the very early stages. At the instant when LH-NL reaches 5 nm (see the first black dot in Fig. S3b), the **TH** has already traveled a distance ≥ 10 nm, indicating that the diffusing **TH** is ahead of the docking LH-NL. Fig. S3b shows a plateau in the dynamics of T338 during which *x*_*T*338_ does not change, while *x_TH_* undergoes large changes. As *x*_*T*338_ fluctuates around 5 nm (between the second and third dots in Fig. S3b), the *x_TH_* diffuses between 4 and 12 nm. The only correlation between *x*_*T*338_ and *x_TH_* seems to be that *x*_*T*338_ set a lower bound for *x*_TH_, meaning *x_TH_* rarely drops below the value of ***x***_*T*338_ by more than ~ 1 nm.

The lack of correlation between the time-dependent changes in ***x_*TH*_*** and ***x***_*T*338_ is more apparent in a plot ***x_*TH*_*** as a function of ***x***_*T*338_ (Fig. S3c). The dotted line corresponds to *x_TH_ = x*_*T*338_. Any data point above (below) the dotted line indicates that **TH** is ahead of (behind) the docking LH-NL. We find that **TH** follows LH-NL up to 3 nm, after which the **TH** is predominately ahead of the LH-NL. At the end of the step, the **TH** moves ~ 16 nm while LH-NL moves only ~ 8 nm. Thus, given the **TH** is ahead of the docking NL during the major duration of the step, it is inaccurate to suggest that **TH** is pulled by NL docking towards the (+) end of the MT. The **TH** reaches the **TBS** at the (+) end through diffusion, with NL docking providing the needed restriction and bias for the **TH** to reach the TBS.

### NL docking is not necessary for detachment of the **TH** from the MT

The mutation simulations also allow us to test the hypothesis that NL has to dock to release the **TH** from the MT. If this hypothesis is correct, we expect that the **TH** would stay bound to the **MT** in the mutation simulations (Fig. 5b). However, **TH** detaches from the **MT** (Fig. 5a) within 5μs (Fig. 5c), in 79% of trajectories. Thus, the intramolecular strain within a two-head-bound state alone is sufficient to detach the TH. When the intramolecular strain is released by deleting the LH, the **TH** stays bound to the **MT** within our simulation window (50μs), and can resists external force up to 3 pN. Therefore, NL docking in the LH is not necessary to release the ADP-bound **TH** from the MT.

**Fig. 5.**
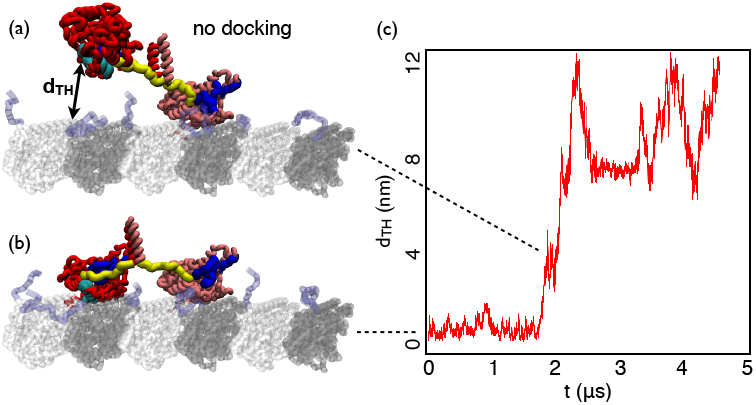
Neck linker docking is not necessary to detach to the **TH** from the **MT**: (a) A snapshot showing **TH** detaches from the MT, while the NL of the LH remains undocked. (b) The initial conformation of the kinesin motor heads in the mutant simulation, where NL docking does not occur. (c) The distance between the **TH** and the **MT** as a function of time recorded in the mutant simulation.

Our observation is not inconsistent with the leading head gating model (37, 38). According to the model, ATP cannot bind to the LH until the **TH** detaches from the MT. In other words, intramolecular stain in the two-head-bound state inhibits ATP binding and thus NL docking to the LH. Therefore, the leading head gate model predicts that **TH** detachment from the **MT** occurs before NL docking to the LH. This prediction is supported by our finding that **TH** detachment from the **MT** does not depend on docking of NL to the LH.

## Discussion

We used simulations of a coarse-grained model with hydrodynamic interactions of the entire MT-Kin complex to (i) quantify the contribution of NL docking and diffusion to the kinesin step, and (ii) illustrate the mechanism by which kinesin avoids side steps. Our model, which is currently the only one to include the effects of **MT** explicitly, is calibrated to reproduce the experimentally measured stall force (10, 11, 39), and the force required to dislodge the motor head from the **MT** (31, 40) (Fig. S2). Interestingly, these two energy scales suffice to capture all the salient features of the kinesin step. Surprisingly, we find that nearly three quarters of the 16 nm step involves almost random search for the **TBS** by the **TH** (Fig. 1 and Fig. 3). The time the **TH** spends in the stochastic search accounts for more than 95% of the duration for completing the 16 nm step (Fig. 1e). However, in order to stay predominantly on the same protofilament of the MT, NL docking is necessary to constrain the diffusion of the **TH**, thus minimizing the probability of side steps (Fig. 4).

Our simulations underscore the importance of large scale diffusive motion within the kinesin step (Fig. 1). One might argue that the stochasticity observed in the simulation is due to the simplicity of the SOP model, which is based on short ranged native contacts. The SOP model allows for fluctuations in the necklinker, before it becomes fully docked to the LH. So could a simulation, where necklinker docking in the LH is triggered by less flexible Go-like model, produce a 16 nm step with decreased stochasticity? It is unlikely for two reasons. First, even assuming that the LH neck linker docks deterministically, **TH** neck linker, which is undocked and connects **TH** and LH, would introduce stochasticity due to the translation of the TH. At the same time, **TH** is free to rotate, regardless of the conformational state of the LH neck linker. Second, temporally neck linker docking accounts for less than 5% of the stepping time. Spatially neck linker docking could pull **TH** towards the plus end by 8 nm at most. The rest of the step has to occur through diffusion, which is independent of the dynamics of neck linker docking. It is also worth noting that Hyeon and Onuchic have shown that the SOP model and the one including dihedral angle potentials in the Go-like model lead to qualitatively similar equilibrium properties, further justifying our used of the SOP model (see the SI in (23)). Thus, the physics of kinesin stepping illustrated here is not dependent on the apparent increase in the flexibility of the model.

It is difficult to capture the bidirectional motion of the kinesin motor domain using only the current experimental techniques, due to the small size of the kinesin motor domain (41) (~ 2nm), and the transient nature of the 16 nm step (11) (~ 30*μ*s). From a theoretical perspective, however, it seems natural to suggest that diffusion must play a key role in kinesin motility. The rate of diffusion (~ 4*μ*s^-1^, for 16 nm) is much faster than the rate of ATP turnover (~ ms^-1^). To experimentally test our prediction of the extent (~ 12nm) and duration (~ 40*μ*s) of diffusion within the kinesin step, would require tracking the motion of the kinesin motor head using optical trap (42) or FIONA (5) at ~ microsecond resolution.

In a recent experiment (22) the motion of the unbound head was tracked high temporal (≈ 55*μ*s) resolution using dark-field microscopy. By tracking the position of a 40nm gold particle connected to the motor head through a biotin-streptavidin construct it was shown that the motion of the unbound head towards the **TBS** occurs by diffusion, in accord with our findings (24). The large hydrodynamic drag due to the gold particle slows down the actual time scale for stepping, suggesting that the extent of diffusion could be even greater than hinted at by these insightful experiments. Even with this high temporal resolution, subtle aspects of search for the **TBS** such as nearly unhindered rotation of the unbound head and multiple attempts to achieve correct orientation of the motor head with respect to the MT, as discovered here, will have to await future experiments involving labeling at multiple sites and higher temporal resolution.

The observation that NL docking is responsible for only ~ 4nm out of the 16 nm step (Fig. 3a) shows that NL docking cannot directly pull the motor head to the next binding site. In this respect, NL docking in kinesin is different from lever arm rotation in myosin. This difference is also supported by experiments (28, 43). If a step is largely due to the conformational change in a mechanical motif (such as docking of the NL or rotation of the lever arm) of a motor protein, then extension of the mechanical motif should lead to an increase of step size and the motor speed. Indeed, the sliding velocity of the myosin motor increases linearly with the length of its lever arm (44). In contrast, extending the NL by stiff helices or double-stranded DNA does not lead to any increase of kinesin speed (28, 43). Therefore, NL docking and lever arm rotation must contribute to motor motility in different ways.

The finding that NL docking in the LH is not necessary for detachment of the **TH** (Fig. 5) is consistent with experiments (38, 45). Using single molecule FRET, Mori ***et.al***. showed that at low ATP concentrations kinesin waits in a one-head-bound state (45). Given that NL docking is triggered by ATP binding, this result suggests that the **TH** can detach spontaneously in absence of NL docking. Further, using fluorescence polarization microscopy, Asenjo ***et.al***. provided evidence that the **TH** is mobile while kinesin waits for ATP (46). Our findings support a recent experiment demonstrating that intramolecular strain generated by NL docking is not necessary to accelerate the detachment of the **TH** (38). According to our simulations, intramolecular strain generated by the two-head-bound state alone could cause the detachment of the **TH** within 5*μ*s in most instances.

From our study we have inferred a physically reasonable mechanism for avoiding side steps: NL docking limits the access to side binding sites and prevents rebinding to the initial site by constraining **TH** diffusion (Fig. 4 and Fig. 6). Such a mechanism has been proposed earlier in an important computational study (23) and simulation from our group (24). Using a computational model based on the potential of mean force experienced by the **TH**, Hyeon and Onuchic (HO) found that the probability of **TH** taking side steps depends on the rate of neck linker docking to the LH (23). In accord with this finding based on equilibrium simulaions, we showed, using detailed stepping dynamics that neck linker docking reduce the probability of side steps (24). In the current study, we calibrated the energy scales associated with docking, by reproducing experimentally measured stall force. We found docking occurs fast within 1.1 *μ*s, which is smaller than 20 *μ*s, the maximum docking time, estimated from the equilibrium simulations (23), to avoid side steps. Therefore, despite differences in the simulation strategies, both studies show that neck linker docking reduces the probability of side steps.

**Fig. 6.**
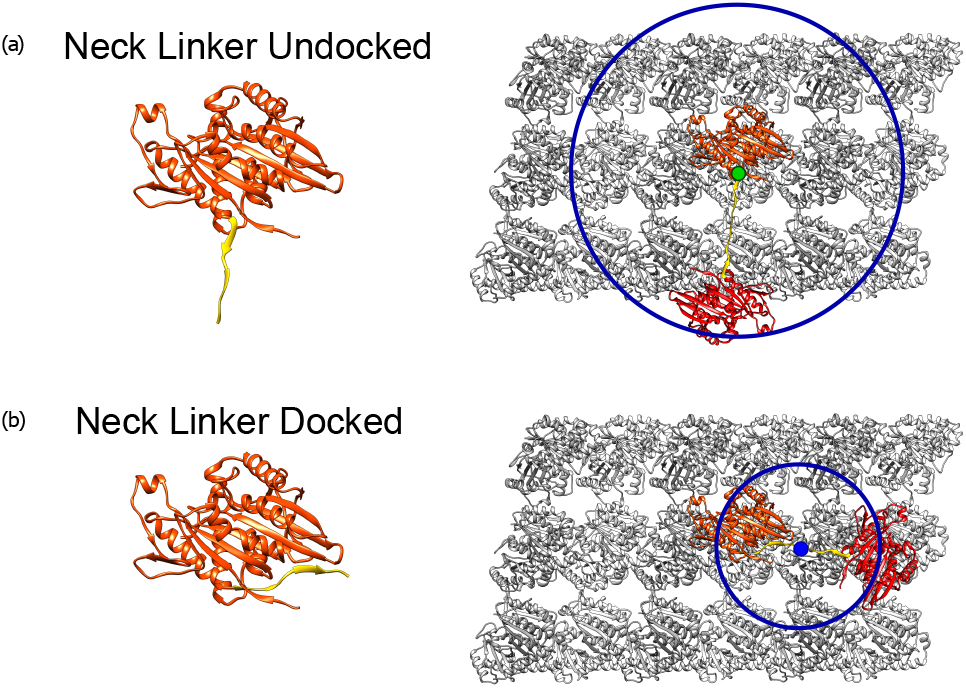
A model for how kinesin chooses its binding site: Schematics of binding sites accessible to the **TH** in the absence (a) and presence (b) of NL docking. The circles illustrate the range of sites that are accessible to a free motor head. In (a) the blue sites can be reached with significant probability if neck linker docking is disfavored. Upon neck linker docking to the LH the most probable site to which the **TH** binds is the one that is along the same protofilament. In both cases the search for various binding sites occurs by stochastic movement after the **TH** detaches from the MT.

However, could side steps be avoided by other mechanisms? For example, instead of limiting the access to side binding sites (Fig. 6b), is it possible to prevent ADP release at these sites (blue dots in Fig. 6a)? Preferential ADP release is supported by previous experiments. ADP-kinesin affinity is lower when NL points towards the minus end of the **MT** (31, 40), implying that the **TH** is more likely to release the bound ADP when it closer to the **TBS** than the IBS. However, the observation that the probability of side steps is larger than 50% for a kinesin mutant with extended NL (28), suggests that **TH** can release ADP at accessible binding sites (blue dots in Fig. 6a) other than the TBS. Therefore, in order to minimize the probability of taking side steps, it is necessary to limit the access of the **TH** to side binding sites by constraining **TH** diffusion (Fig. 6b). One simple and elegant way that nature has solved this problem is by constraining the walk through NL docking.

## Conclusions

Our results unify two seemingly distinct mechanisms in kinesin stepping, the neck linker docking model (8) and the brownian ratchet model (18). Furthermore, we also provide a structural explanation for how kinesin stays on a single microtubule protofilament. Our simulations show that kinesin takes the 16 nm step mainly through Brownian motion, with neck linker docking crucially constraining the Brownian motion of the kinesin motor head.

## Materials and Methods

### Methods

We performed simulations by creating a new coarsegrained (CG) model of the MT-Kin complex (details in the SI). Such models have proved to be efficacious in producing quantitative insights into thestepping kinetics of molecular motors (23, 30, 47—50). Additional details, including the creation of the MT-Kin complex with coiled-coil and cargo based on the Self-Organized Polymer model (51—53), the determination of the two important parameters, and the simulation details are given in the SI. Remarkably, the only two parameters are needed to quantitatively describe the stepping kinetics, with one accounting for the strength of the neck linker attachment to the leading head and the other being the strength of the motor head-MT interaction.

We used Brownian dynamics with hydrodynamic interactions (HIs) to generate large ensemble of stepping trajectories. It is important to point out that in order to observe completion of steps in experimentally relevant time scale HIs must be explicitly included (54). Finally, we note that the use of CG model allows us to generate hundreds of trajectories for both the wild-type and *in silico* mutants, created by assigning zero gain in the energy due to ordering of the neck-linker to the leading head, so that conclusions with sufficient statistics can be drawn.

## ACKNOWLEDGMENTS

We are grateful to Dr. K. H. Downing for providing the coordinates of MT. We appreciate interactions with Prof. Michael E. Fisher during the course of this work, which was done as a partial fulfillment of the doctoral thesis (ZZ) requirement in 2012. We are grateful to Mauro Mugnai and Naoto Hori for constructive suggestions. This work was supported by a grant from the National Science Foundation (CHE 16-61946). Additional support from the Collie-Welch Reagents Chair (F-0019) is gratefully acknowledged.

## Notes

Authors declare no conflict of interest.

## References

1. Howard J (2001) Mechanics of Motor Proteins and the Cytoskeleton. (Sinauer Associates, Sunderland, MA).

2. Vale RD (2003) The molecular motor toolbox for intracellular transport. Cell 112:467–480.

3. Block SM (2007) Kinesin motor mechanics: Binding, stepping, tracking, gating, and limping. Biophys. J. 92:2986–2995.

4. Hancock WO (2016) The Kinesin-1 Chemomechanical Cycle: Stepping Toward a Consensus. Biophys. J. 110(6):1216–1225.

5. Yildiz A, Tomishige M, Vale RD, Selvin PR (2004) Kinesin walks hand-over-hand. Science 303:676–678.

6. Asbury CL, Fehr AN, Block SM (2003) Kinesin moves by an asymmetric hand-over-hand mechanism. Science 302:2130–2134.

7. Hua W, Chung J, Gelles J (2002) Distinguishing inchworm and hand-over-hand processive kinesin movement by neck rotation measurements. Science 295:844–848.

8. Rice S, et al. (1999) A structural change in the kinesin motor protein that drives motility. Nature 402:778–784.

9. Schnitzer M, Block S (1997) Kinesin hydrolyses one ATP per 8-nm step. Nature 388(6640):386–390.

10. Svoboda K, Block S (1994) Force and velocity measured for single kinesin molecules. Cell 77:773–784.

11. Carter NJ, Cross RA (2005) Mechanics of the kinesin step. Nature 435:308–312.

12. Spudich JA, Sivaramakrishnan S (2010) Myosin VI: an innovative motor that challenged the swinging lever arm hypothesis. Nat. Rev. Mol. Cell Biol. 11:128–137.

13. Gennerich A, Carter AP, Reck-Peterson SL, Vale RD (2007) Force-induced bidirectional stepping of cytoplasmic dynein. Cell 131:952–965.

14. Reck-Peterson SL, et al. (2006) Single-molecule analysis of dynein processivity and stepping behavior. Cell 126:335–348.

15. Gennerich A, Vale RD (2009) Walking the walk: how kinesin and dynein coordinate their steps. Curr. Opin. Cell Biol. 21(1):59–67.

16. Schief WR, Howard J (2001) Conformational changes during kinesin motility. Curr. Opin. Cell Biol. 13(1):19–28.

17. Carter NJ, Cross RA (2006) Kinesin’s moonwalk. Curr. Opin. Cell Biol. 18(1):61–67.

18. Taniguchi Y, Nishiyama M, Ishii Y, Yanagida T (2005) Entropy rectifies the Brownian steps of kinesin. Nat. Chem. Biol. 1:342–347.

19. Tomishige M, Stuurman N, Vale RD (2006) Single-molecule observations of neck linker conformational changes in the kinesin motor protein. Nat. Struct. Mol. Biol. 13:887–894.

20. Asenjo AB, Weinberg Y, Sosa H (2006) Nucleotide binding and hydrolysis induces a disorder-order transition in the kinesin neck-linker region. Nat. Struct. Mol. Biol. 13:648–654.

21. Khalil AS, et al. (2008) Kinesin’s cover-neck bundle folds forward to generate force. Proc. Nat. Acad. Sci. USA 105:19247–19252.

22. Isojima H, Lino R, Aiitani Y, Noji H, Tomishige M (2016) Direct observation of intermediate states during the stepping motion of kinesin-1. Nat. Chem. Biol. 12:290–297.

23. Hyeon C, Onuchic JN (2007) Mechanical control of the directional stepping dynamics of the kinesin motor. Proc. Natl. Acad. Sci. U. S. A. 104(44):17382–17387.

24. Zhang Z, Thirumalai D (2012) Dissecting the Kinematics of the Kinesin Step. Structure 20:628–640.

25. Kutys ML, Fricks J, Hancock WO (2010) Monte Carlo Analysis of Neck Linker Extension in Kinesin Molecular Motors. PLOS COMP. BIOL. 6(11):e1000980.

26. Ray S, Meyhofer E, Milligan RA, Howard J (1993) Kinesin follows the microtubules protofilament axis. J. Cell Biol. 121:1083–1093.

27. Block SM, Asbury CL, Shaevitz JW, Lang MJ (2003) Probing the kinesin reaction cycle with a 2D optical force clamp. Proc. Natl. Acad. Sci. U. S. A. 100:2351–2356.

28. Yildiz A, Tomishige M, Gennerich A, Vale RD (2008) Intramolecular strain coordinates kinesin stepping behavior along microtubules. Cell 134:1030–1041.

29. Svoboda K, Schmidt CF, Schnapp BJ, Block SM (1993) Direct observation of kinesin stepping by optical trapping interferometry. Nature 365:721–727.

30. Hyeon C, Thirumalai D (2011) Capturing the essence of folding and functions of biomolecules using coarse-grained models. Nat. Comm. 2:487.

31. Uemura S, et al. (2002) Kinesin-microtubule binding depends on both nucleotide state and loading direction. Proc. Natl. Acad. Sci. USA 99:5977–5981.

32. Woehlke G, et al. (1997) Microtubule interaction site of the kinesin motor. Cell 90(2):207–216.

33. Nitta R, Okada Y, Hirokawa N (2008) Structural model for strain-dependent microtubule activation of Mg-ADP release from kinesin. Nat. Struct. Mol. Biol. 15:1067–1075.

34. Hinczewski M, Tehver R, Thirumalai D (2013) Design principles governing the motility of myosin V. Proc. Natl. Acad. Sci. 110(43):E4059–E4068.

35. Purcell T, Morris C, Spudich J, Sweeney H (2002) Role of the lever arm in the processive stepping of myosin V. Proc. Natl. Acad. Sci. 99(22):14159–14164.

36. Kodera N, Yamamoto D, Ishikawa R, Ando T (2010) Video imaging of walking myosin V by high-speed atomic force microscopy. Nature 468(7320):72–77.

37. Guydosh N, Block S (2006) Backsteps induced by nucleotide analogs suggest the front head of kinesin is gated by strain. Proc. Natl. Acad. Sci. 103(21):8054–8059.

38. Andreasson JOL, et al. (2015) Examining kinesin processivity within a general gating framework. ELIFE 4:e07403.

39. Visscher K, Schnitzer M, Block S (1999) Single kinesin molecules studied with a molecular force clamp. Nature 400(6740):184–189.

40. Uemura S, Ishiwata S (2003) Loading direction regulates the affinity of ADP for kinesin. Nat. Struct. Biol. 10:308–311.

41. Kull F, Sablin E, Lau R, Fletterick R, Vale R (1996) Crystal structure of the kinesin motor domain reveals a structural similarity to myosin. Nature 380:550–555.

42. Guydosh NR, Block SM (2009) Direct observation of the binding state of the kinesin head to the microtubule. Nature 461(7260):125–128.

43. Miyazono Y, Hayashi M, Karagiannis P, Harada Y, Tadakuma H (2010) Strain through the neck linker ensures processive runs: a DNA-kinesin hybrid nanomachine study. EMBO J. 29:93–106.

44. Uyeda T, Abramson P, Spudich J (1996) The neck region of the myosin motor domain acts as a lever arm to generate movement. Proc. Natl. acad. Sci. 93(9):4459–4464.

45. Mori T, Vale RD, Tomishige M (2007) How kinesin waits between steps. Nature 450:750–U15.

46. Asenjo AB, Sosa H (2009) A mobile kinesin-head intermediate during the ATP-waiting state. Proc. Natl. Acad. Sci. U. S. A. 106:5657–5662.

47. Hyeon C, Onuchic JN (2007) Internal strain regulates the nucleotide binding site of the kinesin leading head. Proc. Natl. Acad. Sci. U. S. A. 104(7):2175–2180.

48. Hwang W, Lang MJ, Karplus M (2008) Force generation in kinesin hinges on cover-neck bundle formation. Structure 16:62–71.

49. Hariharan V, Hancock WO (2009) Insights into the Mechanical Properties of the Kinesin Neck Linker Domain from Sequence Analysis and Molecular Dynamics Simulations. Cell. Mol. Bioeng. 2:177–189.

50. Grant BJ, et al. (2011) Electrostatically Biased Binding of Kinesin to Microtubules. PLoS. Biol. 9:e1001207.

51. Hyeon C, Dima RI, Thirumalai D (2006) Pathways and kinetic barriers in mechanical unfolding and refolding of RNA and proteins. Structure 14:1633–1645.

52. Hyeon C, Lorimer GH, Thirumalai D (2006) Dynamics of allosteric transitions in GroEL. Proc. Natl. Acad. Sci. USA 103:18939–18944.

53. Chen J, Dima RI, Thirumalai D (2007) Allosteric communication in dihydrofolate reductase: Signaling network and pathways for closed to occluded transition and back. J. Mol. Biol. 374:250–266.

54. Goldtzvik Y, Zhang Z, Thirumalai D (2016) Importance of Hydrodynamic Interactions in the Stepping Kinetics of Kinesin. J. Phys. Chem. B. 120(8):2071–2075.

